# Thermal cycling hyperthermia sensitizes non-small cell lung cancer A549 cells to EGFR-tyrosine kinase inhibitor Erlotinib

**DOI:** 10.1101/2022.05.03.490542

**Authors:** Guan-Bo Lin, Wei-Ting Chen, Yu-Yi Kuo, Hsu-Hsiang Liu, You-Ming Chen, Shr-Jeng Leu, Chih-Yu Chao

## Abstract

Molecular targeted therapy has emerged as a mainstream treatment for non-small cell lung cancer (NSCLC), the most common lung cancer, which has been the leading cause of cancer death in both genders. Erlotinib (Erl), a targeted therapy inhibiting epidermal growth factor receptor (EGFR) pathways, has been proved to have noticeable response rate for NSCLC cells. However, limited efficacy arises due to intrinsic and acquired resistance among most NSCLC patients. Therefore, sensitizers are required to potentiate the efficacy of Erl in NSCLC treatment. The study proposed a novel thermal therapy, thermal cycling hyperthermia (TC-HT), as a supplement to amplify the effects of Erl. We found that TC-HT remarkably reduced the half-maximal inhibitory concentration (IC50) of Erl to as little as 0.5 μM and demonstrated that TC-HT could sensitize A549 NSCLC cells to Erl via the downstream of EGFR signaling cascades. Furthermore, combination treatment induced G2/M cell cycle arrest, and inhibition of cell proliferation and migration. Besides, via raising the high temperature of TC-HT slightly, TC-HT treatment alone can produce excellent antineoplastic effect without hurting the normal cells. The method is expected to be applicable to other combination therapies and may be a starter for more sophisticated, side-effect-free anticancer treatments.

## 1 Introdution

As one of the most common type of cancer, lung cancer was the largest source of cancer-related deaths in 2020, according to World Health Organization (WHO). In particular, Non-small cell lung cancer (NSCLC) accounts for 85% of lung cancer deaths and there has yet to be effective treatment for it (1). Available therapies for NSCLC treatment include surgery, chemotherapy, radiation, and targeted therapy. As chemotherapy alone is not effective, due to low response rate, there is urgent need for combination treatment or new agents targeting one or more molecular pathways.

Targeted treatment, meant to inhibit cancer malignancies with focus on critical molecules, has been regarded as a promising therapy to treat NSCLC patients (2). Among various targeted therapies, anti-epidermal growth factor receptor (EGFR) has been the one with most extensive application in clinical treatment of NSCLC (3). EGFR is a receptor tyrosine kinase of the ErbB family which plays a key role in cell proliferation, survival, differentiation, and migration (4). When activated by ligand binding, EGFR will autophosphorylate and trigger downstream signaling cascades, including mitogen-activated protein kinase (MAPK) and phosphatidylinositol 3-kinase (PI3K)-Akt pathways (5), leading to cell cycle progression and inhibiting pro-apoptotic proteins. Therefore, several EGFR-inhibiting agents, such as monoclonal antibodies and tyrosine kinase inhibitors (TKIs) have been developed for anticancer treatments (6). Erlotinib (Erl), a first generation EGFR-TKI, has exhibited significantly better response rate and benefits in NSCLC patients than conventional platinum-based chemotherapy (7), capable of prolonging the survival of patients with advanced NSCLC after chemotherapy. Regrettably, most tumors will develop acquired resistance to Erl over time (8), plus side effects such as rash and diarrhea for long-term administration of high-dose Erl (9). Therefore, it is worthwhile to evaluate the effect of combination treatment in potentiating the efficacy of Erl at low dose, in place of chemotherapy drugs, such as cisplatin and pemetrexed, which have been applied as sensitizers to enhance the efficacy of Erl (10, 11), with side effects such as hypertension and severe diarrhea (12).

In this work, we proposed a novel thermal therapy, thermal cycling hyperthermia (TC-HT), as a substitute for chemotherapy drugs or agents, to investigate whether such physical cue, TC-HT, may be an effective sensitizer to amplify the anticancer effect of Erl. Given the safety and effectiveness of TC-HT reported in our previous studies (13, 14), we aimed to probe the effect of combining Erl and TC-HT in A549 NSCLC cell line. The results showed that TC-HT significantly enhanced the anticancer effect of Erl by sensitizing A549 cells to Erl. As a consequence, the Erl dosage required to achieve significant anticancer effect was greatly reduced. The approach is expected to solve the problems of low response rate and acquired resistance for cancer drugs in NSCLC patients, while reducing side effects significantly. The enhanced anticancer effect was achieved in part via EGFR and downstream pathways as shown by Western blot analysis and flow cytometry. The results show that Erl and TC-HT synergistically weaken the phosphorylation of EGFR and modulate the phosphorylated c-Jun N-terminal kinases (p-JNK) and Akt protein expressions, thereby inducing the mitochondria-dependent apoptosis. Besides, combination treatment of Erl and TC-HT regulated phosphorylated p38 (p-p38) and cell division cycle protein 2 (Cdc2), thereby inducing G2/M cell cycle arrest. Meanwhile, the results of the colony formation assay and wound healing assay indicated that the combination treatment has significant inhibitory effect on cell growth and migration. Noteworthily, TC-HT alone, to our surprise, can exhibit potent anticancer effect by raising the high temperature setting slightly, which is comparable to the in vitro efficacy of chemotherapy drugs. Moreover, normal lung cell line IMR-90 was almost unharmed under the same TC-HT treatment condition.

In this paper, the study demonstrated for the first time that TC-HT can potentiate the anticancer efficacy of Erl on A549 cells, thereby reducing their needed dosage in treatment. Moreover, by raising the high temperature of TC-HT slightly, TC-HT treatment could hinder growth of A549 cancer cells, without drug administration. These findings suggest that combination of TC-HT and Erl is a promising approach in anticancer treatment, which sheds light on combining TC-HT with other anticancer drugs for potential treatment.

## 2 Materials and methods

### 2.1 Cell culture

The human NSCLC cell line A549 and normal lung cell line IMR-90 were purchased from the Bioresource Collection and Research Center of the Food Industry Research and Development Institute (BCRC, Hsinchu, Taiwan). Both cell lines were seeded on cell culture flasks and cultured in DMEM (HyClone; Cytiva, Marlborough, MA, USA) supplemented with 10% fetal bovine serum (HyClone; Cytiva) and 1% penicillin-streptomycin (Gibco; Thermo Fisher Scientific, Inc., Waltham, MA, USA) in a humidified incubator with 5% CO_2_ at 37°C.

### 2.2 Drug treatment and TC-HT exposure

Erl, purchased from MedChemExpress (MCE, Monmouth Junction, NJ, USA), was dissolved in dimethyl sulfoxide (DMSO) (Sigma-Aldrich; Merck KGaA, Darmstadt, Germany) to a concentration of 5 mM as the stock solution and stored at −20°C. A549 cells were seeded in 96-well plates overnight before treated with various concentrations of Erl. The TC-HT groups were exposed to a thermal cycling treatment with a high temperature period for 3 min and a cooling period for 30 sec, and this protocol was repeated continuously for ten cycles, as shown in Figure 1A. The actual temperatures sensed by the cells at the bottom of the well were measured by a needle thermocouple. For the combination treatment, TC-HT was applied for 1 h before Erl treatment. After the single or combination treatment, the cells were maintained in the cell culture incubator for an additional 72 h for further analyses.

**Figure 1.**
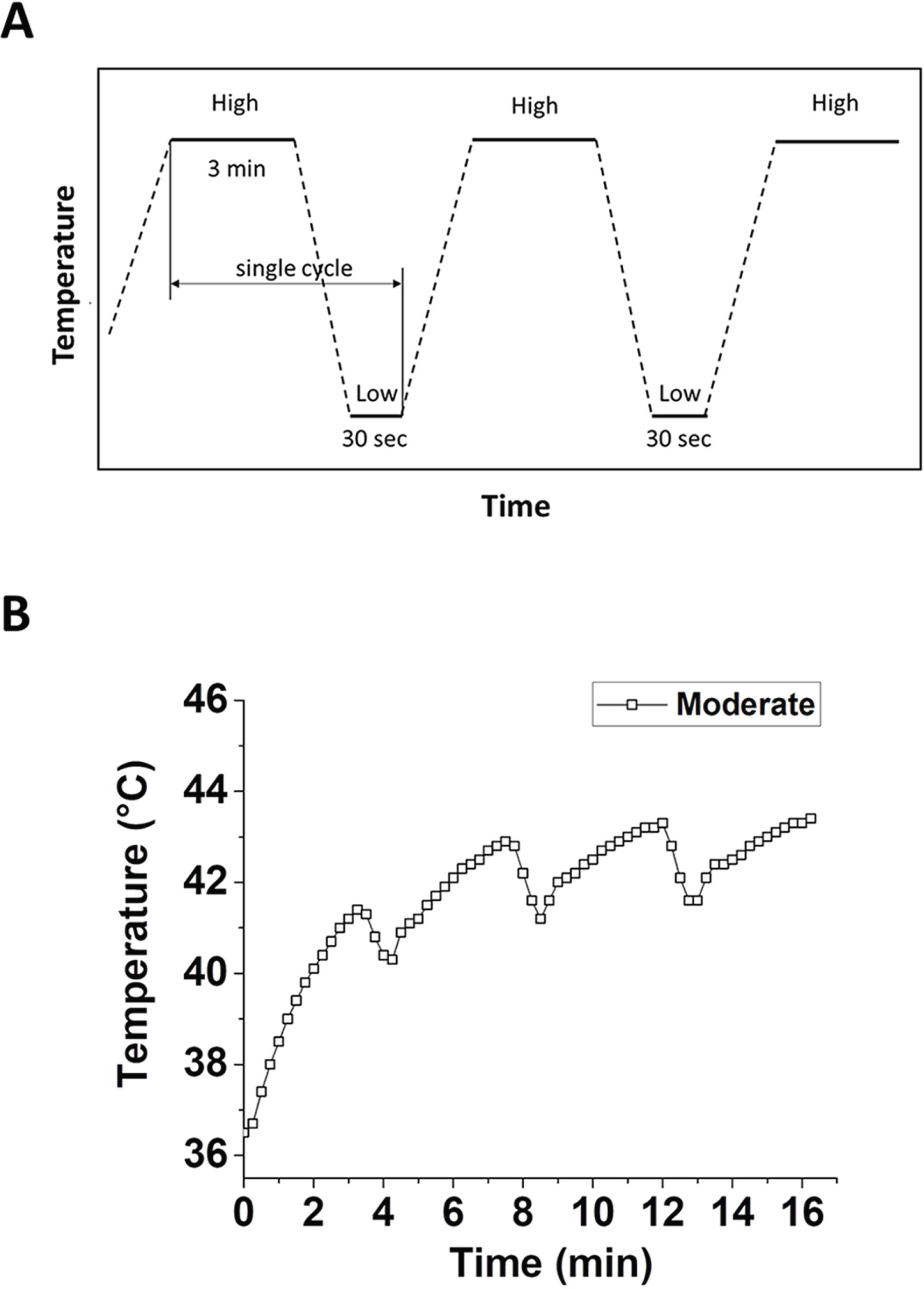
In vitro-applied TC-HT. **(A)** Schematic representation of the high and low temperatures and their duration settings during TC-HT treatment. **(B)** Actual temperature with moderate temperature TC-HT setting in the culture well measured every 15 sec by a needle thermocouple.

### 2.3 Cell viability assay and morphology

The viability of A549 cells was assessed by 3-(4,5-dimethylthiazol-2-yl)-2,5-diphenyltetrazolium bromide (MTT) (Sigma-Aldrich; Merck KGaA) assay. After drug or TC-HT treatment, the medium was replaced by DMEM containing 0.5 mg/mL MTT, and incubated for 4 h at 37°C. The formazan crystals were dissolved by 10% sodium dodecyl sulfate (SDS) (Bioman Scientific Co., Ltd., Taipei, Taiwan) overnight. Thereafter, the optical density of formazan solution was measured at 570 nm using Multiskan GO spectrophotometer (Thermo Scientific, Hudson, NH). Background absorbance at 690 nm was subtracted, and the final absorbance value was expressed as a percentage of the untreated controls to represent the cell viability. Cell morphology of A549 cells under different treatment was observed and photographed by the Zyla 5.5 sCMOS camera (Andor, Belfast, UK).

### 2.4 Western blot analysis

After treatment with Erl or TC-HT or in combination, cells were washed with ice-cold phosphate buffered saline (PBS) and lysed in RIPA lysis buffer (EMD Millipore, Billerica, MA, USA) on ice for 30 min. Cell lysates were clarified by centrifugation at 23,000 × g for 30 min at 4°C, and then the protein concentration in the supernatant fraction was measured by the Bradford protein assay (Bioshop, Burlington, Canada). Proteins (20-40 μg) were subjected to 10% SDS-PAGE and electrotransferred to polyvinylidene fluoride (PVDF) membranes (EMD Millipore). After incubation in a blocking buffer (5% bovine serum albumin/TBST) for 1 h at room temperature, the membranes were immunoblotted with diluted primary antibodies at 4°C overnight. The specific primary antibodies against phosphorylated EGFR (p-EGFR), poly ADP-ribose polymerase (PARP), p-JNK, phosphorylated Akt (p-Akt), total Akt (t-Akt), MutT homolog 1 (MTH1) (Cell Signaling Technology, Inc., Danvers, MA, USA), p-p38, Cdc2 and glyceraldehyde-3-phosphate dehydrogenase (GAPDH) (GeneTex, Inc., Irvine, CA, USA) were used. After being washed with TBST, the membranes were incubated with horseradish peroxidase-conjugated secondary antibodies (Jackson ImmunoResearch Laboratories, Inc., West Grove, PA, USA). Images were visualized with an enhanced chemiluminescence substrate (Advansta, Inc., Menlo Park, CA, USA) and quantified by Amersham Imager 600 imaging system (AI600, GE Healthcare Life Sciences, Chicago, IL, USA).

### 2.5 Mitochondrial membrane potential measurement

After treatment with Erl or TC-HT or in combination, cells were washed and resuspended with PBS followed by staining with 20 nM 3,3’-dihexyloxacarbocyanine iodide (DiOC_6_(3)) (Enzo Life Sciences International Inc., Farmingdale, NY, USA) for 30 min at 37°C in the dark. The fraction of mitochondrial depolarization in cells is indicated by the decrease of fluorescence intensity measured by a flow cytometer (FACSVerse; BD Biosciences, San Jose, CA, USA).

### 2.6 Wound healing assay

A549 cells were seeded and cultured as a confluent monolayer in 35 mm diameter Petri dishes. In vitro, wounds were made by scratching straight lines across the cell monolayer using a pipette tip (10 µ L). Non-adherent cells and debris were removed by gently rinsing the cells with PBS (HyClone; Cytiva). After PBS was discarded and replaced with fresh medium, the cells were treated with Erl or TC-HT or in combination. Each wound was observed and photographed by the Zyla 5.5 sCMOS camera (Andor) at 0 h and 24 h after treatments. The distances between wound edges were measured and analyzed using ImageJ software.

### 2.7 Colony formation assay

A549 cells were seeded at 300 cells/dish in 35 mm Petri dishes overnight for cell adherence. Subsequently, cells were treated with Erl or TC-HT or in combination. Cell medium was replaced 24 h after the treatment, and the cells were cultured for additional 10 days, with fresh medium replaced every 3 days during the culture period. At last, the cells were fixed with 4% paraformaldehyde (PFA) (Sigma-Aldrich; Merck KGaA) for 10 min and stained with 0.1% crystal violet (Sigma-Aldrich; Merck KGaA) to make them visible and counted. The number of colonies in each group was normalized to control group.

### 2.8 Cell cycle analysis

After 24 h treatment with 10 μM Erl or TC-HT or in combination, the cells were harvested, rinsed with PBS, and fixed with 70% ethanol at 4°C for 30 min before staining. The cells were then stained for 30 min in the dark with propidium iodide (PI) and ribonuclease A (RNase A) (Gibco; Thermo Fisher Scientific, Inc.). Subsequently, the stained cells were subject to the cell cycle analysis by using a flow cytometer (FACSVerse; BD Biosciences), and data were analyzed with FlowJo software.

### 2.9 Statistical analysis

The statistical results were expressed as the average from at least three independent experiments and the error bars are ± standard deviation. Statistical analyses were performed by using OriginPro 2015 software (OriginLab). Differences of statistical significance were determined by one-way analysis of variance (ANOVA) followed by Tukey’s post-hoc test. P < 0.05 was considered to indicate a statistically significant difference.

## 3 Results

### 3.1 TC-HT for in vitro application

The thermal cycling treatment was applied with a modified PCR machine as described previously (13, 14). The schematic temperature and duration settings are shown in Figure 1A, where the temperature was increased to the desired high temperature setting and maintained for 3 min, followed by a natural cooling period for 30 sec. In practice, the heating device was switched off in the cooling process, and the low temperature setting was chosen to be 37°C to mimic human body temperature. A single cycle containing a high temperature and a low temperature period was repeated for 10 times in this study. Figure 1B showed the actual temperature in the culture well measured by a thermocouple at moderate temperature TC-HT setting. As can be seen in Figure 1B, the cycling temperature comes to an equilibrium state after the third heating period, thereafter the temperature cycling between 41.5∼43°C for moderate temperature TC-HT. In this work, we adopted the moderate temperature TC-HT treatment (41.5∼43°C) in the following experiments to study the synergistic anticancer effect of Erl and TC-HT in A549 cancer cells.

### 3.2 TC-HT potentiates the anticancer efficacy of Erl in A549 NSCLC cells

To determine the effect of combination Erl and TC-HT treatment on human NSCLC cell viability, A549 cells were treated with different concentrations of Erl with or without TC-HT. At 72 h after the treatments, cell viability was assessed by MTT assay. As shown in Figure 2A, Erl exhibited antineoplastic effect in a concentration-dependent manner. The half-maximal inhibitory concentration (IC50) of Erl was about 10 µM in A549 cells after 72 h treatment. Compared to the treatment with Erl alone, TC-HT therapy sensitizes A549 cells to Erl and synergistically reduces the cell viability to about 30% of the control group (Figure 2A). It is noteworthy that the combination of Erl and TC-HT demonstrated synergistic ability as compared with Erl monotherapy, which remarkably reduced the IC50 to as little as 0.5 µM (up to 20-fold decrease). To verify the effect of this combined treatment in normal cells, the normal lung cell line IMR-90 was used and applied with the same treatment condition. As shown in Figure 2B, the cell viability of IMR-90 normal lung cell line was almost unaffected by Erl and/or TC-HT regardless of the treatment procedure. Additionally, as shown in Figure 2C, Erl and TC-HT combination treatment resulted in notable morphological changes in A549 cells, including shrinkage and fragmentation of cells as well as a decreased number of cells. The result implies that TC-HT is not only a riskless sensitizer to natural compounds (13, 14), but also an effective and safe sensitizer to increase the anticancer effect of targeted therapy drug.

**Figure 2.**
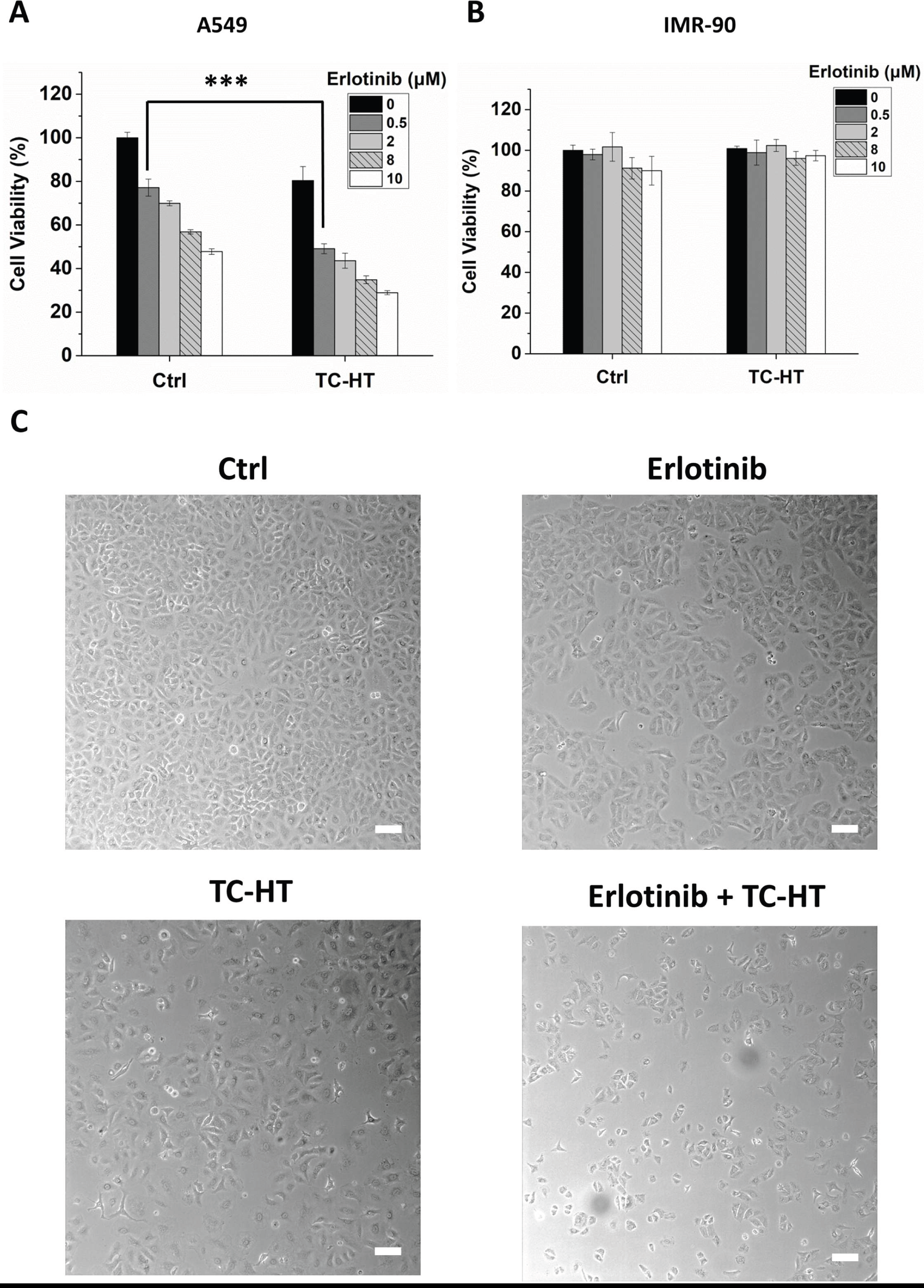
Effect of Erl or TC-HT or in combination on the cell viability and morphological changes of A549 cells. **(A)** MTT viability assay of A549 NSCLC cells treated with different concentrations of Erl or in combination with moderate temperature TC-HT treatment (41.5∼43°C). **(B)** MTT viability assay of IMR-90 normal lung cells treated with 5 and 10 μM Erl or in combination with moderate temperature TC-HT treatment (41.5∼43°C). **(C)** Representative images of morphological changes of A549 cells under the light microscope after treatment with 10 μM Erl, moderate temperature TC-HT treatment (41.5∼43°C) or in combination. Scale bar = 100 μm. Data represent the mean ± standard deviation (n=3). Statistical significance was determined by ANOVA followed by Tukey’s post-hoc test (\*\*\**P* < 0.001).

### 3.3 Effect of Erl combined with TC-HT on EGFR protein expression in A549 cells

Here we examined the signaling pathways involved in the anticancer mechanism of Erl and TC-HT combination treatment. With NSCLC cells frequently exhibiting EGFR overexpression (15), targeted therapy drugs were applied to inhibit the activity. Meant specifically for cancers with EGFR mutation, Erl inhibits cancer cells by blocking the intracellular phosphorylation of tyrosine kinase with such mutation (7). However, NSCLC may develop acquired resistance to Erl over time, according to some studies (8). In the study, as shown in Figure 3, Erl alone caused a slight decrease in EGFR phosphorylation compared to the control cells. Moreover, the study finds that while TC-HT alone can significantly inhibit p-EGFR expression, the combination of Erl and TC-HT can induce more pronounced inhibition effect on p-EGFR expression. It suggests that TC-HT may modulate the resistance of A549 cells to Erl and alter the activity of EGFR downstream pathways.

**Figure 3.**
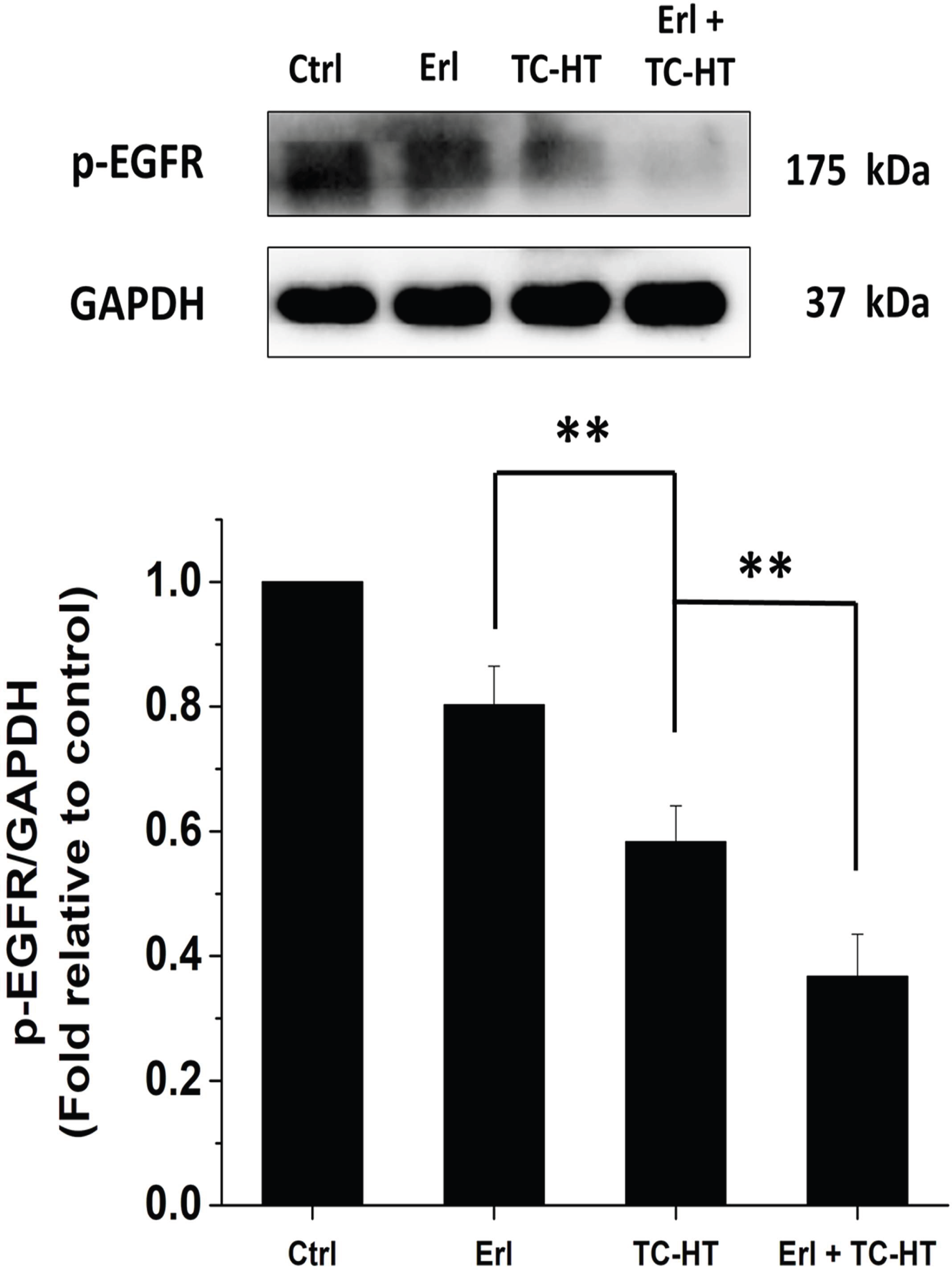
Effect of Erl combined with TC-HT on p-EGFR protein expression in A549 cells. Western blot analysis of p-EGFR protein expression in A549 cells treated with 10 μM Erl, moderate temperature TC-HT, and the combination treatment. The expression levels of p-EGFR were normalized to GAPDH. Each relative expression level was compared with control and represented as fold relative to control. Data represent the mean ± standard deviation (n=3). Differences of statistical significance were determined by one-way ANOVA followed by Tukey’s post-hoc test (\*\**P* < 0.01 and \**P* < 0.05).

### 3.4 Effect of Erl combined with TC-HT on EGFR signaling pathways in A549 cells

In the study, we then examined the expressions of JNK and Akt proteins (16) in A549 cells. JNK protein, an EGFR downstream protein belonging to the MAPK family, is deemed to be responsive to stress stimuli and heat shock, with its phosphorylation capable of altering the activities of many proteins that reside in mitochondria or act in the nucleus and trigger apoptosis (17). As shown in Figure 4A, TC-HT treatment significantly increased the expression of p-JNK protein, compared with the control or Erl-treated A549 cells. The result showed that TC-HT alone can effectively promote apoptosis. Furthermore, we found that the treatment combining Erl and TC-HT is much more effective in upregulating the p-JNK expression than TC-HT alone, as it can boost apoptosis even more. On the other hand, another EGFR downstream protein Akt is involved in cellular survival pathways by inhibiting apoptotic process (18). Once activated by phosphorylation, Akt actively regulates a number of transcription factors facilitating the expression of survival proteins. Figure 4B showed that the expression of phosphorylated Akt (p-Akt) in Erl-treated A549 cells is lower than the untreated cells, while the inhibitory effect on phosphorylation of Akt increases further in TC-HT treatment and is even higher in the combination Erl and TC-HT treatment. Meanwhile, total Akt (t-Akt) protein exhibits a trend similar to that of p-Akt protein, suggesting that decrease in the phosphorylation of Akt in these treatments is caused by the reduction in the amount of Akt, leading to a weakened survival signal. Moreover, it is known that both JNK and Akt signaling pathways are important regulators in influencing mitochondrial function. In addition, the mitochondria-dependent apoptosis is associated with a loss in mitochondrial membrane potential (MMP), causing the release of cytochrome c and other apoptotic factors (19). To confirm that the apoptosis caused by Erl and TC-HT could be related to mitochondrial disruption, we assessed MMP with DiOC6(3) fluorescence staining via flow cytometry. As shown in Supplementary Figure S1, we confirmed that Erl and TC-HT individually had no effect on MMP level compared to control, but the combination treatment caused significant MMP depolarization in A549 cells, indicating mitochondrial dysfunction in their apoptosis mechanism.

**Figure 4.**
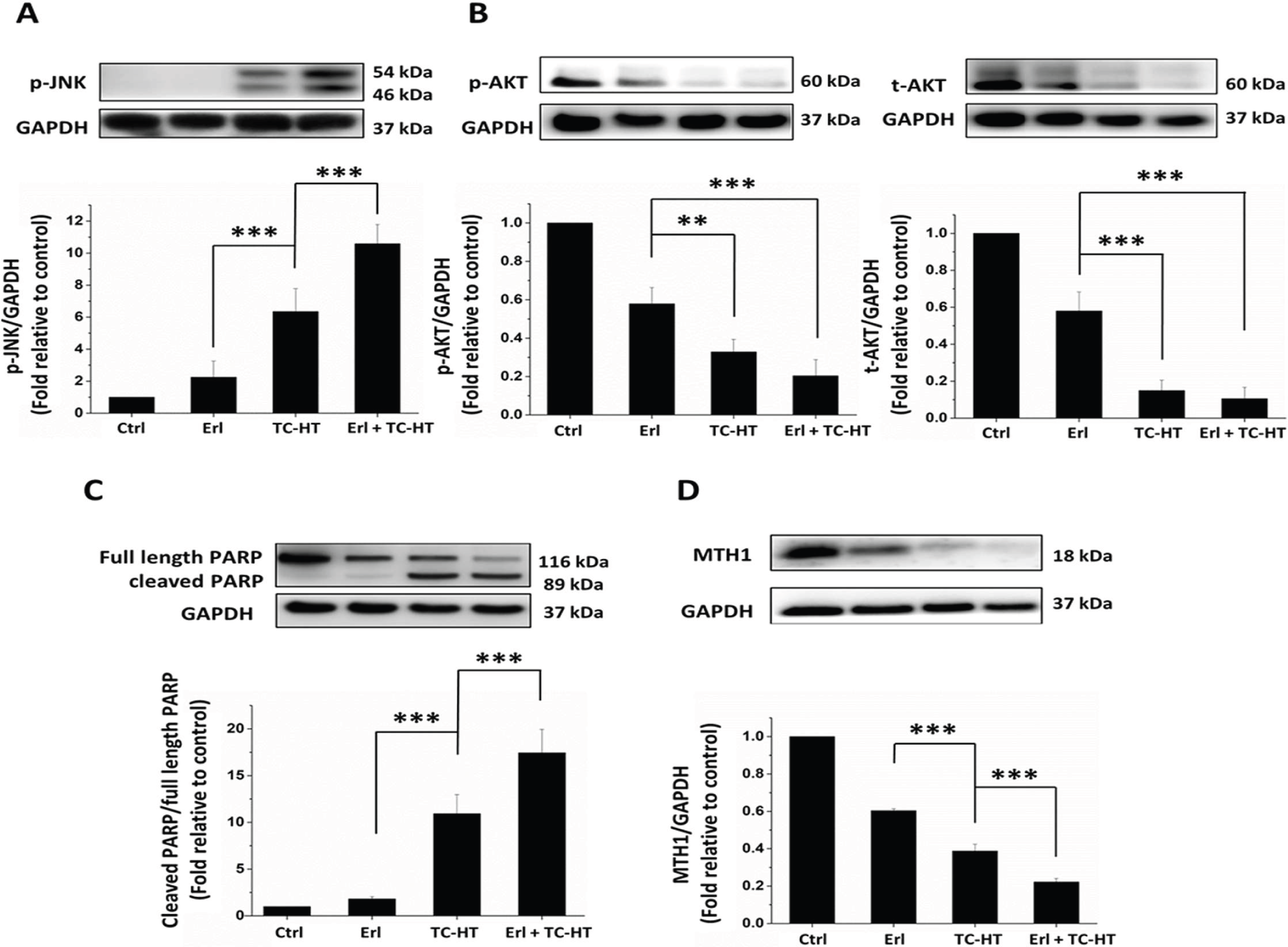
Effect of Erl combined with TC-HT treatment on survival- and apoptosis-related protein expressions in A549 cells. The anticancer experiments were conducted in A549 cancer cells treated with 10 μM Erl, moderate temperature TC-HT, and the combination treatment. Western blot analysis of p-JNK **(A)**, p-Akt and t-Akt **(B)**, cleaved PARP and full length PARP **(C)**, MTH1 **(D)** protein expressions. The expression levels of p-JNK, p-Akt, t-Akt, and MTH1 were normalized to GAPDH and cleaved PARP was normalized to full length PARP. Each relative expression level was compared with control and represented as fold relative to control. Data represent the mean ± standard deviation (n=3). Differences of statistical significance were determined by one-way ANOVA followed by Tukey’s post-hoc test (\*\*\**P* < 0.001 and \*\**P* < 0.01).

The injured mitochondria releases cytochrome c into the cytoplasm, cleaving caspase 9 and thus activating caspase 3 in the downstream (20), which enters further the nucleus and cleaved PARP. It should be noted that PARP plays an important role in mitochondria-mediated apoptosis, in addition to being a key enzyme for DNA repair. In apoptosis, PARP protein is typically cleaved and inactivated, thereby suppressing DNA repair and causing programmed cell death (21). To understand the apoptosis mechanism in A549 cells triggered by the combination Erl and TC-HT treatment, the study evaluated the expression of apoptosis-related proteins with Western blot analysis. Figure 4C showed that the ratio of cleaved PARP to full length PARP in TC-HT-treated cells was much higher than that in Erl-treated cells. Moreover, we find that the ratio of cleaved PARP to full length PARP in combination Erl and TC-HT treatment was much stronger than that in TC-HT treatment alone, indicating the combination treatment enhances apoptosis of A549 cells via mitochondrial pathway. In the same vein, MTH1 protein has received increasing attention in cancer treatment, due to its key role in DNA repair (22). High expression of MTH1 is deemed to be a sign for the metastasis of NSCLC (23). Therefore, the study examined MTH1 protein expression via Western blot analysis. As shown in Figure 4D, MTH1 expression was found to decrease in Erl-treated cells, while the inhibitive effect on MTH1 protein expression was much higher in TC-HT treatment, which increased further in the combination Erl and TC-HT treatment. The result reveals that the combination of Erl and TC-HT may prevent the MTH1-related DNA repair and induce killing of cancer cells via apoptosis.

### 3.5 The combination treatment of Erl and TC-HT causes G2/M cell cycle arrest in A549 cells

To further evaluate the anticancer effects of combination Erl and TC-HT treatment on human NSCLC, the cell cycle progression in the A549 cells was examined by flow cytometric analysis. As shown in Figures 5A and 5B, treatment with moderate TC-HT led to a higher extent (22.95±0.56%) on the cell cycle arrest in the G2/M phase compared to the group treated with Erl (13.06±1.16%) and the untreated cells (12.66±1.18%). Meanwhile, the combination of Erl and moderate TC-HT caused significant accumulation of cells in the G2/M phase (33.77±2.31%) with a concomitant reduction of cells in the G0/G1 phase. Besides, no significant differences in the S phase were observed among the various treatments. In the following, we examined the relevant proteins involved in the G2/M progression to investigate the mechanism of the combined treatment effect on cell cycle distribution. Cdc2 protein is a core regulator in cell cycle, and Cdc2 binds to cyclin B complex guiding G2/M cell cycle transition (24, 25). It was reported that the inhibition of Cdc2 expression resulted in G2/M cell cycle arrest (26–28). In the study, as shown in Figure 5C, the expression of Cdc2 reduced significantly in combination Erl and TC-HT treatment. Meanwhile, as a well-known MAPK member, p38 plays an important role in cell cycle regulation, and many studies have shown that activation of p38 pathway can induce G2/M cell cycle arrest via Cdc2 suppression in human NSCLC cells (29–31). Consistently, we found that the expression of p-p38 was markedly enhanced under the combination treatment of Erl and moderate TC-HT (Figure 5D). Taken together, these results suggested that the combined treatment could synergistically induce cell cycle arrest in the G2/M phase by increasing the activation of p38 while reducing Cdc2 expression in cells, ultimately inhibiting the proliferation of human lung cancer A549 cells.

**Figure 5.**
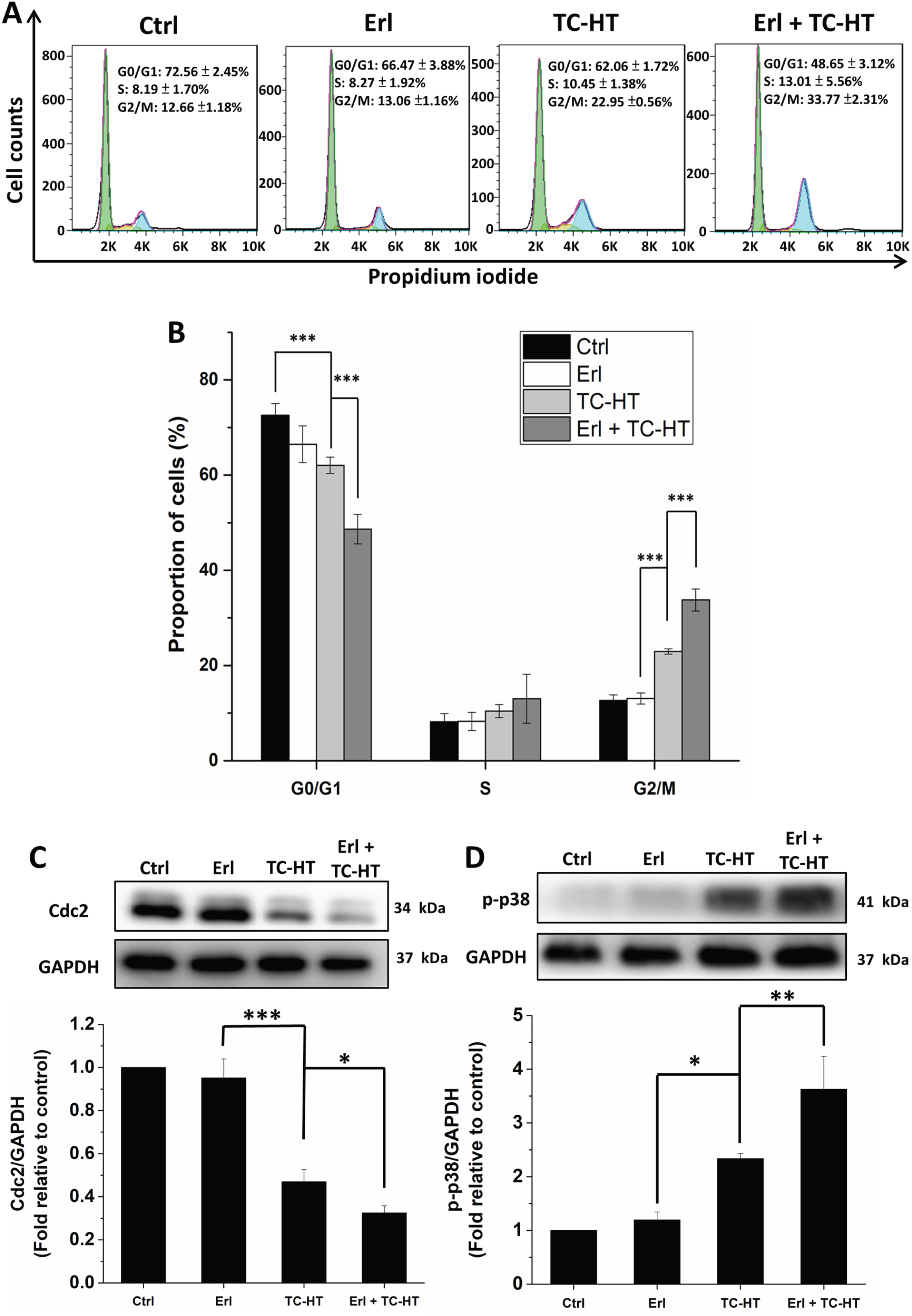
Effect of Erl or TC-HT or in combination on the G2/M cell cycle arrest and protein expressions of Cdc2 and p-p38 in A549 cells. The experiments of cell cycle and protein analysis were conducted in A549 cancer cells treated with 10 μM Erl, moderate temperature TC-HT (41.5∼43°C), and the combination treatment. **(A)** Flow cytometry analysis on cellular DNA content profiles of each group of A549 cells. **(B)** The proportion of A549 cells in G0/G1, S and G2/M phases of the indicated group. Western bolt analysis of Cdc2 **(C)** and p-p38 **(D)** protein expressions. The expression levels of p-p38 and Cdc2 were normalized to GAPDH. Each relative expression level was compared with control and represented as fold relative to control. Data represent the mean ± standard deviation (n=3). Statistical significance was determined by one-way ANOVA followed by Tukey’s post-hoc test (\*\*\**P* < 0.001, \*\**P* < 0.01, and \**P* < 0.05).

### 3.6 Anti-proliferation and anti-migration effects of the combination treatment (Erl + moderate temperature TC-HT) in A549 NSCLC cells

Cancer mortality is associated with cancer recurrence and metastasis, both of which are common clinical problems in NSCLC (32, 33). Therefore, it is very important to reduce the growth and migration ability of cancer cells. To further confirm the anti-proliferative and anti-migration activities of Erl and TC-HT combination treatment in NSCLC, we performed wound healing assay and colony formation assay on A549 cells. As shown in Figures 6A and 6C, Erl-treated cells migrated over a smaller area than control cells, while cells treated with TC-HT alone exhibited more reduction in migration. Furthermore, the Erl and TC-HT combination treatment caused more significant suppression of A549 cell migration compared to those of the single treatments. In comparison to the control group, the wound closure rate in the Erl-treated group and the TC-HT-treated group was 83.8% and 59.2%, respectively, while the combined treatment’s wound closure rate was only 35.9%. The results indicated that the migration ability of A549 cells was significantly suppressed after the application of Erl and TC-HT combination treatment. On the other hand, we found that colony formation in A549 cells treated with Erl or TC-HT was decreased, whereas the colony formation in cells treated with combination treatment was further and greatly reduced (Figures 6B and 6D). Overall, the combination treatment has the ability to impede the A549 cell growth and migration, indicating the potential anticancer efficacy of Erl and TC-HT combination treatment.

**Figure 6.**
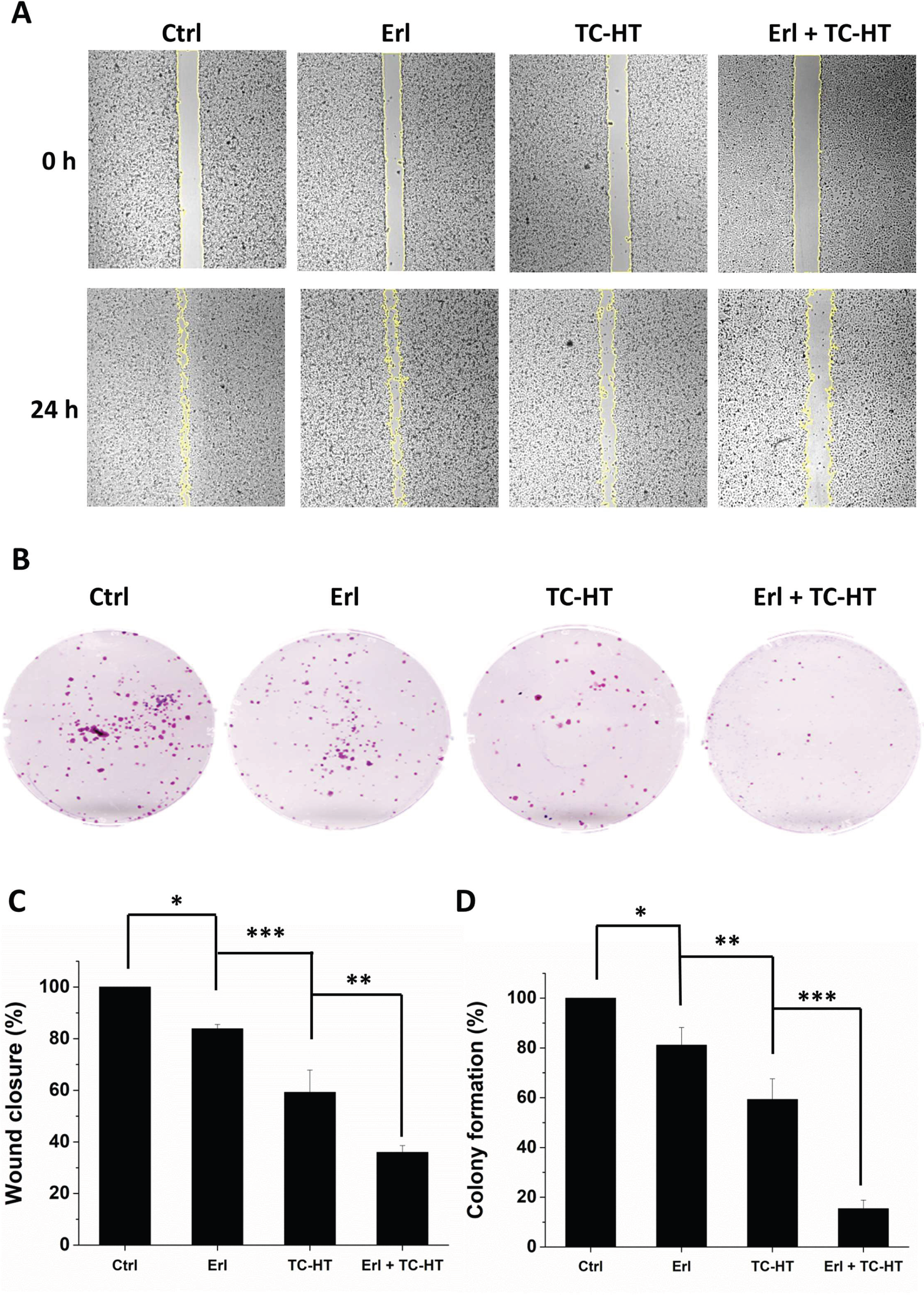
Effect of Erl or TC-HT or in combination on the inhibition of A549 cell colony formation and migration. **(A)** Wound healing assay to determine the effect of Erl, TC-HT or combination treatment on the migration ability of A549 cells. After scratch gaps were made, A549 cells were treated with 10 µM Erl, moderate temperature TC-HT, and the combination treatment, respectively. **(B)** The results of colony formation assay for A549 cells treated with 10 µM Erl, moderate temperature TC-HT, and the combination treatment, respectively. **(C)** Wound closure rate for A549 cells was determined as the percentage of the area closed after 24 h from the initial wound area, and each group was normalized to control group and expressed as a fraction of 100. The areas of cell-free gaps were measured and quantified using ImageJ. **(D)** Colony formation rate for A549 cells at 10 days after treatment with 10 µM Erl, moderate temperature TC-HT, and the combination treatment, respectively. Each group was normalized to control group and represented in the form of fractions of 100. The colony counting was performed using ImageJ. Differences of statistical significance were determined by one-way ANOVA followed by Tukey’s post-hoc test (\*\*\**P* < 0.001, \*\**P* < 0.01, and \**P* < 0.05).

## 4 Discussion

The study demonstrated the synergistic anticancer effect of the combination of Erl (an EGFR-tyrosine kinase inhibitor) and TC-HT on A549 NSCLC cells. EGFR is a receptor tyrosine kinase critical for the initiation and development of malignant tumors via MAPK and PI3K/Akt pathways. Although the EGFR-TKIs such as Erl are meant to target and inhibit the EGFR pathways, their clinical benefit has been limited, due to the intrinsic and acquired resistance to TKIs among most NSCLC patients. Amid the efforts to combat the resistance of EGFR-TKIs (34), combination treatment has been deemed the most promising approach to overcome the resistance problem (35). Several studies have investigated the effect of combining Erl and other anticancer agents such as cisplatin, monoclonal antibodies, aspirin, and capsaicin (10, 36–38), aiming to enhance the therapeutic effect. However, the combined use of multiple drugs in clinical applications is often hampered by unpredictable drug interactions or side effects. Therefore, we look for alternate way to enhance the anticancer effect by combining physical stimulation with the use of drug. The study demonstrated for the first time the effect of TC-HT physical stimulation in boosting the anticancer effect of Erl. In comparison to Erl or TC-HT mono treatment, the combination treatment produced a higher inhibitory effect on A549 NSCLC cells, without hurting the normal lung cells (Figure 2), thereby circumventing drug interaction problem and significantly reducing the needed dosage of Erl, which could lessen side effects. To elucidate the molecular mechanisms underlying the anticancer effect of the combination treatment on A549 NSCLC cells, the expressions of proteins in the apoptotic pathway were evaluated. We observed that TC-HT significantly potentiated the efficacy of the targeted therapy drug Erl in inhibiting the phosphorylation of EGFR (Figure 3), which amplified the suppressing effect of Erl on A549 NSCLC cells. Subsequently, we investigated the JNK and Akt proteins in the EGFR downstream pathway. The activation of JNK is recognized to induce apoptosis (17), while the activation of Akt serves to inhibit apoptosis (18). Additionally, both JNK and Akt proteins play a crucial role in the regulation of mitochondrial function [19], and thus investigating the activation status of these proteins contributes to a better understanding of cellular apoptotic tendency. As illustrated in Figures 4A and 4B, our results showed that the combination of TC-HT and Erl increased p-JNK expression while decreasing p-Akt expression. Concurrently, the combination treatment of Erl and TC-HT significantly increased MMP depolarization in A549 cells (Supplementary Figure S1). In addition, the cleavage of PARP protein and the downregulation of MTH1 expression are both noteworthy targets for contributing apoptosis in cancer cells (21–22). In this study, we find that a significant increase in the expression of cleaved PARP is accompanied by a substantial decrease in MTH1 expression (Figures 4C and 4D). These findings suggest that the combination treatment of Erl and TC-HT effectively enhances the apoptosis of A549 NSCLC cells. The cell cycle dysregulation, a common feature of cancer, could be ameliorated through the modulation of cell cycle regulatory proteins, offering the potential to attenuate the uncontrolled proliferation of cancer cells. In this study, we found that the combination treatment of Erl and TC-HT resulted in G2/M phase arrest (Figure 5A and 5B). Cdc2 protein is considered to be a core regulator involved in the G2/M cell cycle transition (24, 25). As shown in Figure 5C, significant downregulation of Cdc2 expression was found in the combination treatment group of Erl and TC-HT. Furthermore, the p38 protein was worth investigating, as previous studies in A549 NSCLC cells demonstrated that increased p38 phosphorylation resulted in reducing the level of Cdc2 expression, eventually leading to G2/M cell cycle arrest (29–31). In this study, we also found that the combined treatment of Erl and TC-HT significantly increased the expression of p-p38. These results indicated that the combination treatment of Erl and TC-HT effectively reduced the occurrence of mitosis in A549 NSCLC cells, representing the inhibition of cancer cell proliferation. Previous studies have suggested that Erl alone has only a mild or even negligible effect on G2/M arrest, while combining Erl with other approaches holds promise for improving its efficacy (39, 40). Similarly, our investigation also found a weak influence of Erl on G2/M arrest when Erl was administered alone, whereas a notable enhancement of G2/M arrest was observed when Erl was combined with TC-HT. It is worth noting that TC-HT, as a physical stimulation, offers a new way to avoid potential drug-drug interactions, meriting further investigation in medical applications.

Inhibiting cancer cell proliferation, recurrence, and migration has long been considered an important part of cancer treatment, and high rates of recurrence and metastasis are also prevalent concerns in NSCLC (32, 33). In this study, we observed that the combined treatment of Erl and TC-HT effectively reduced the cell viability of A549 NSCLC cells (Figure 2), and also induced the G2/M cell cycle arrest (Figure 5). Additionally, we investigated the proliferation and migration of A549 cancer cells under the combined treatment of Erl and TC-HT. The results in Figures 6A and 6C highlight significant findings concerning the migratory behavior of A549 cells under different treatments. We find that single Erl or TC-HT treatment has a reduced migration area in comparison to control cells. Notably, the combination treatment of Erl and TC-HT is more effective in inhibiting A549 NSCLC cells migration than the single Erl or TC-HT treatment. Moreover, A549 NSCLC cells treated with Erl or TC-HT displayed reduced colony formation, whereas the combination treatment of Erl and TC-HT exerted a more profound inhibitory effect on A549 cells (Figures 6B and 6D) than the single treatments. Collectively, these findings illuminate the combined treatment of Erl and TC-HT as a promising anticancer strategy, significantly impeding both A549 NSCLC cell migration and colony formation, suggesting its potential as a prospective anticancer treatment.

Hyperthermia has long been a promising cancer treatment method, to be applied alone or in combination with other conventional therapies (41). The treatment, however, often causes cell damage, due to overdose, a practical problem which can now be overcome by TC-HT, thanks to its ability to control thermal dosage, as its intermittent cooling process can avoid excessive thermal dosage accumulation and subsequent cytotoxic cell damage. Excessive thermal dosage may induce mitochondrial damage and oxidative stress not only in cancer cells but also in normal cells (42). However, studies have shown that cancer cells have higher ROS levels and thus they are more sensitive to thermal stress (43). In this study, the results show that moderate temperature TC-HT treatment (41.5∼43°C) alone inhibited the cell viability of A549 NSCLC cells slightly, reducing their viability to 80% of the control group, without harming IMR-90 normal lung cells (Figures 2A and 2B). To further examine the effect of TC-HT alone on cancer cells at higher temperature, the temperature setting in TC-HT application was raised to the range of high temperature TC-HT treatment (42.5∼45.6°C), as shown in 7A. We found that the high temperature TC-HT treatment alone resulted in significant inhibition of the cell viability of A549 NSCLC cells, with their viability dropping to only 17.9% of that of the control group, as shown in Figure 7B. It is noteworthy that under high temperature TC-HT treatment (42.5∼45.6°C), the additional incorporation of Erl in the range of 0-10 μM did not cause a significant additional decrease in cell viability. Moreover, the TC-HT treatment at high temperature (42.5∼45.6°C) alone did not have much influence on the viability of IMR-90 normal lung cells, which remained at 80% of that of the control group, even in the case of combined treatment with 10 μM Erl (Figure 7C). The results suggest that TC-HT is a potential anticancer treatment for NSCLC and perhaps other cancer cells, capable of reducing the reliance on drugs, and further research to explore its efficacy is under progress.

**Figure 7.**
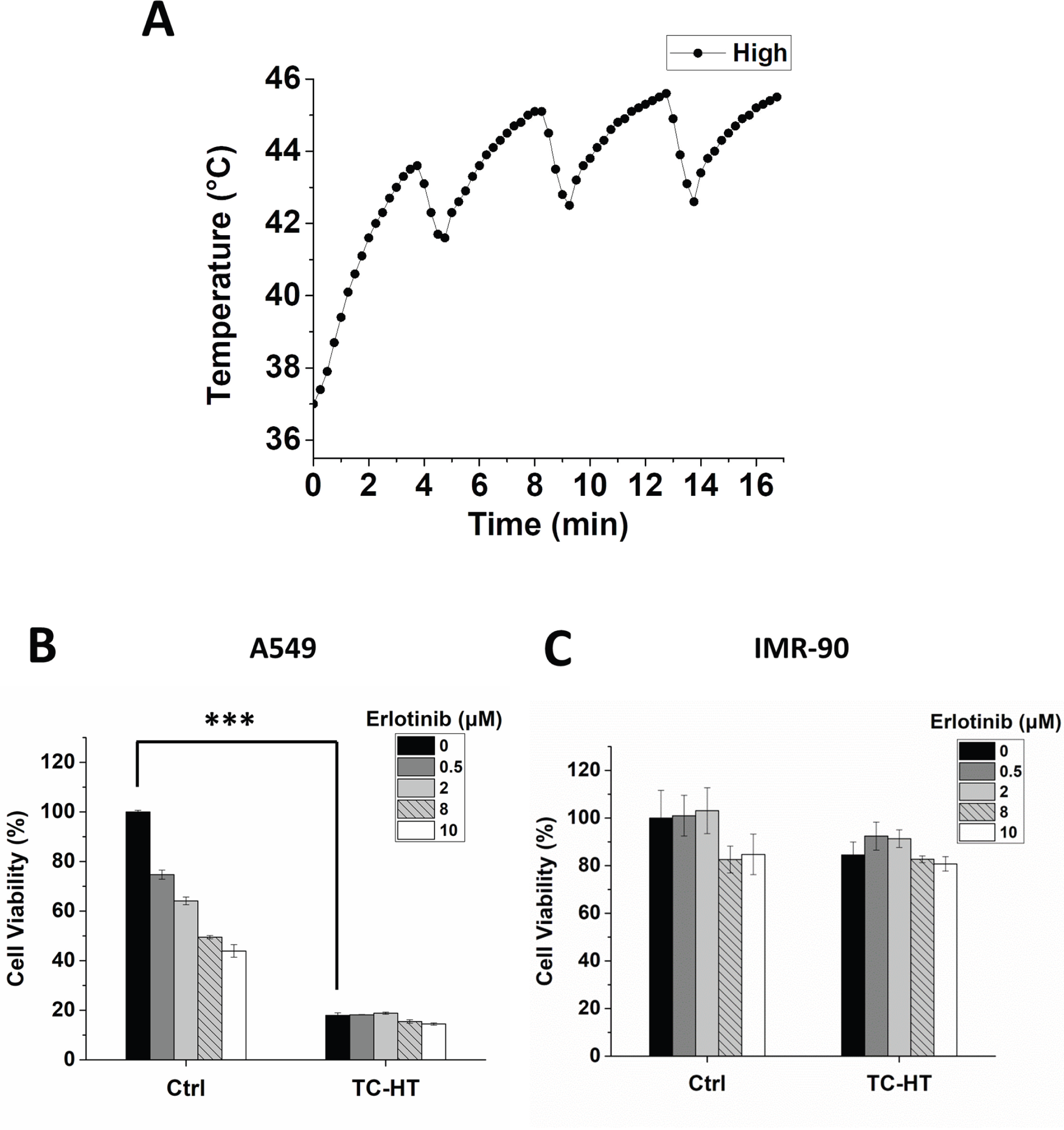
Effect of high temperature TC-HT or Erl or in combination on the cell viability. **(A)** Actual temperature with high temperature TC-HT setting in the culture well measured every 15 sec by a needle thermocouple. **(B)** MTT viability assay of A549 NSCLC cells treated with different concentrations of Erl or in combination with high temperature TC-HT treatment (42.5∼45.6°C). **(C)** MTT viability assay of IMR-90 normal lung cells treated with various concentrations of Erl or in combination with high temperature TC-HT treatment (42.5∼45.6°C). Data represent the mean ± standard deviation (n=3). Statistical significance was determined by one-way ANOVA followed by Tukey’s post-hoc test (\*\*\**P* < 0.001).

In conclusion, the study demonstrates for the first time that TC-HT, a novel thermal treatment, is capable of sensitizing A549 cells to EGFR-TKI Erl, via the downstream of EGFR signaling cascades, including the MAPK and PI3K-Akt pathways. Our results show that TC-HT can enhance the anticancer efficacy of Erl on A549 cells, thereby reducing Erl drug dosage and side effects significantly. Furthermore, TC-HT may become a drug-free cancer treatment method via slightly raising the high temperature of TC-HT in its application. These findings point out the potential for TC-HT in combination therapy with other chemotherapy or targeted therapy drugs, and further studies are needed to examine specific TC-HT parameters in treating different types of cancers.

## Data availability statement

The datasets generated for this study are available on request to the corresponding author.

## Author contributions

C-YC conceived and supervised the study. G-BL, W-TC, and C-YC designed the study. G-BL and W-TC performed experiments and collected data. All authors contributed to data analysis. G-BL, W-TC, and C-YC wrote the manuscript. All authors contributed to the article and approved the final manuscript.

## Acknowledgments

The authors would like to acknowledge the service provided by the Research Core Facilities 3 Laboratory of the Department of Medical Research at National Taiwan University hospital for use of flow cytometry system.

## Supplementary materials

**Supplementary Figure 1.**
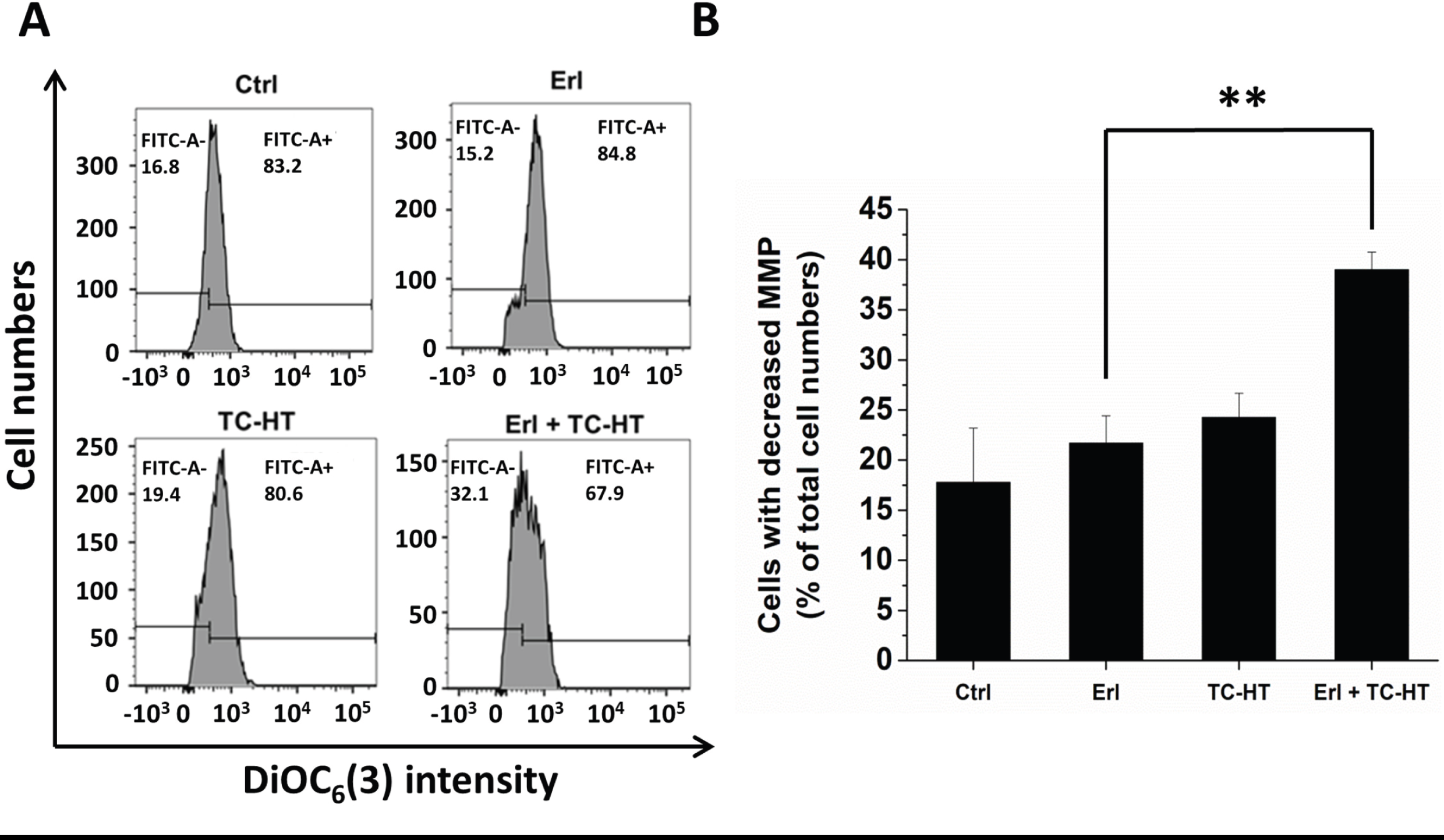
Flow cytometric analysis of MMP in A549 cells treated with Erl and/or TC-HT. A549 cells were treated with 10 μM Erl, moderate temperature TC-HT, and the combination of both treatments. **(A)** MMP was evaluated by the DiOC_6_(3) fluorescent dye and detected by flow cytometer. **(B)** Quantification of the cells with decreased MMP was measured. Data represent the mean ± standard deviation (n=3). Statistical significance was determined by one-way ANOVA followed by Tukey’s post-hoc test (**P < 0.01).

